# Quantitative Deciphering of Mammalian Histamine Receptors through Mathematical Genomics

**DOI:** 10.64898/2026.01.07.698197

**Authors:** Sk. Sarif Hassan, Debaleena Nawn, Pritam Goswami, Nabanita Mukherjee, Moumita Sil, Sujan Roy, Arunava Goswami, Satadal Das, Vladimir N. Uversky

## Abstract

Histamine receptors (HRH1–HRH4) are G-protein coupled receptors (GPCRs) that mediate essential physiological functions, including neurotransmission, gastric acid secretion, immune regulation, and presynaptic autoregulation. Despite their importance, systematic comparative analyses across mammalian histamine receptor sequences remain scarce. In this study, we performed a comprehensive evaluation of HRH1–HRH4 across multiple mammalian species, integrating sequence homology, invariant residue mapping, amino acid substitutions, compositional frequencies, Shannon entropy, polystring distributions, intrinsic disorder profiling, and phylogenetic clustering. HRH2 and HRH1 were highly conserved among primates, HRH3 showed strong cohesion within rodents, while HRH4 exhibited pronounced divergence consistent with immune-related specialization. Invariant residues localized to transmembrane helices and activation motifs (D107, W428/Y431, NPxxY), underscoring strict evolutionary constraints on ligand binding, receptor stability, and G-protein coupling. Substitutions were confined to non-essential lipid-facing and loop regions, predominantly conservative in nature, enabling diversification without disrupting the GPCR fold. Amino acid frequency and entropy analyses revealed hydrophobic dominance with subtype-specific enrichment of polar residues, while disorder profiling identified HRH1 as the most dynamic and HRH2 as the most structurally constrained. Polystring analysis highlighted conserved motifs (WWW, PP) alongside subtype- and species-specific repeats, reflecting evolutionary strategies balancing receptor stability with adaptive flexibility. Phylogenetic clustering confirmed subtype-specific cohesion, with HRH3 and HRH4 forming compact clades, HRH1 showing moderate dispersion, and HRH2 forming the most isolated cluster. Collectively, these findings demonstrate that mammalian histamine receptor evolution is governed by conserved biophysical cores and selective variability, offering insights into structural conservation, functional diversification, and translational relevance for drug design and model selection.

## 1. Introduction

Histamine, first described by Lewis and Grant in 1924 [1, 2], is a fast-acting biogenic amine widely distributed in mammalian tissues [3, 4]. Through the activation of four rhodopsin-like G protein–coupled receptors (H1–H4), histamine regulates diverse physiological and pathological processes, most notably neurotransmission, gastric acid secretion, and inflammation [5, 6, 7, 8, 9, 10]. These receptors, discovered sequentially and named accordingly, exemplify the structural hallmark of GPCRs—seven transmembrane *α*-helices that mediate signaling with ligands such as histamine, acetylcholine, leukotrienes, and adrenaline [11]. GPCR genes constitute approximately 1% of the human genome and underpin essential functions ranging from development and vision to cognition and respiration [12].

GPCRs are commonly classified using the A–F system based on sequence similarity and conserved features of their seven transmembrane domains. This scheme defines six receptor classes across vertebrates and invertebrates. Class A (rhodopsin-like receptors) is the largest group, comprising about 80% of GPCRs, and includes receptors for hormones, neurotransmitters, and light. These receptors share a conserved structure of seven transmembrane helices with an additional cytoplasmic helix near the C-terminus [13]. GPCRs signal through heterotrimeric G proteins, where the inactive G*α* subunit binds GDP [14]. Upon ligand binding, G*α* dissociates from G*βγ* and exchanges GDP for GTP, activating down-stream effectors that regulate signaling, gene expression, and cellular structure [15]. GTP hydrolysis and re-association with G*βγ* restore the basal state, enabling rapid cycling between active and inactive forms. Although G*βγ* can initiate independent pathways, GPCR specificity is primarily determined by the G*α* subunit, encoded by 20 genes grouped into four families: G*α*i, G*α*s, G*α*q, and G*α*12/13 [16]. Histamine receptors couple to distinct G-protein pathways: HRH1 (487 amino acids) couples to Gq, activating phospholipase C and elevating intracellular calcium and inositol phosphate, thereby mediating vascular permeability, inflammation, bronchoconstriction, and allergic responses (Figure 1) [17]. HRH2 (359 amino acids) couples to G*α*s, stimulating cAMP production and regulating gastric acid secretion, antibody synthesis, nerve transmission, and muscle function (Figure 1) [17]. HRH3 (445 amino acids) couples to Gsi/o proteins, modulating cAMP, calcium signaling, MAPK activity, and ion channels; predominantly presynaptic, it influences neurotransmitter release and neuroendocrine signaling [18]. HRH4 (390 amino acids), also Gsi/o-coupled, is enriched in immune tissues and regulates immune responses, T-cell balance, and mast and dendritic cell activity [19].

**Figure 1:**
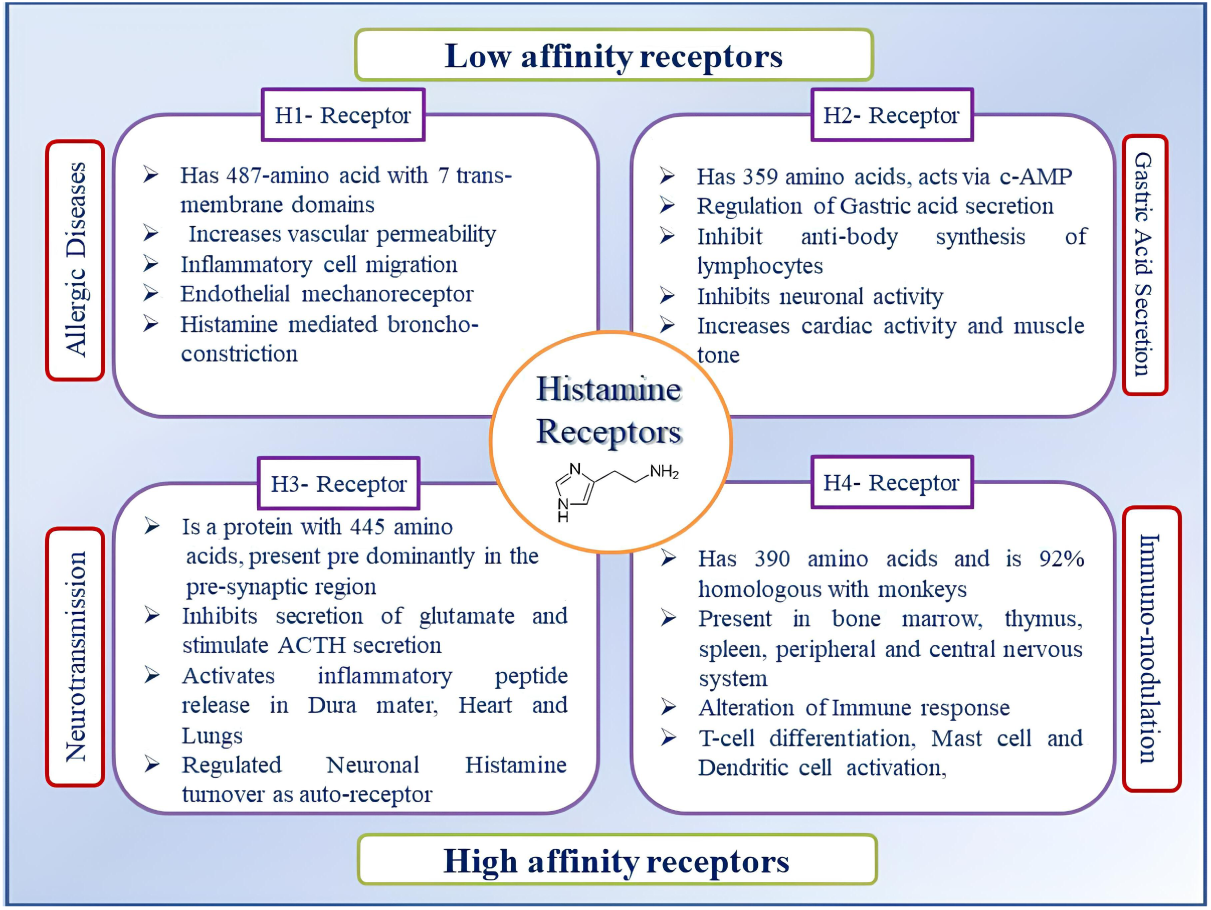
Histamine receptor distribution and their primary identities

**Figure 2:**
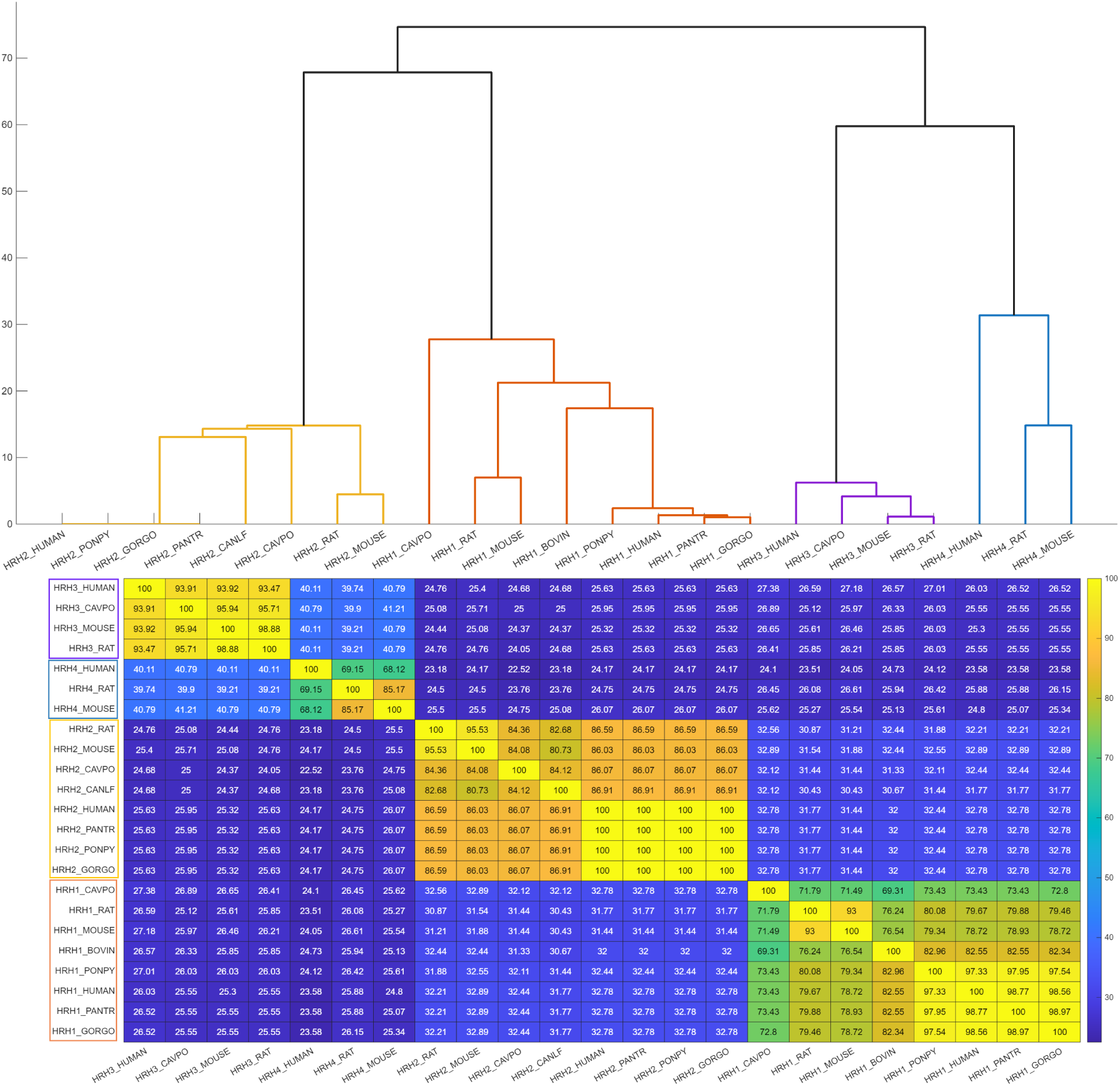
(A) Dendrogram of histamine receptors of all four types; (B) Heatmap matrix of similarity percentages across all histamine receptors.

**Figure 3:**
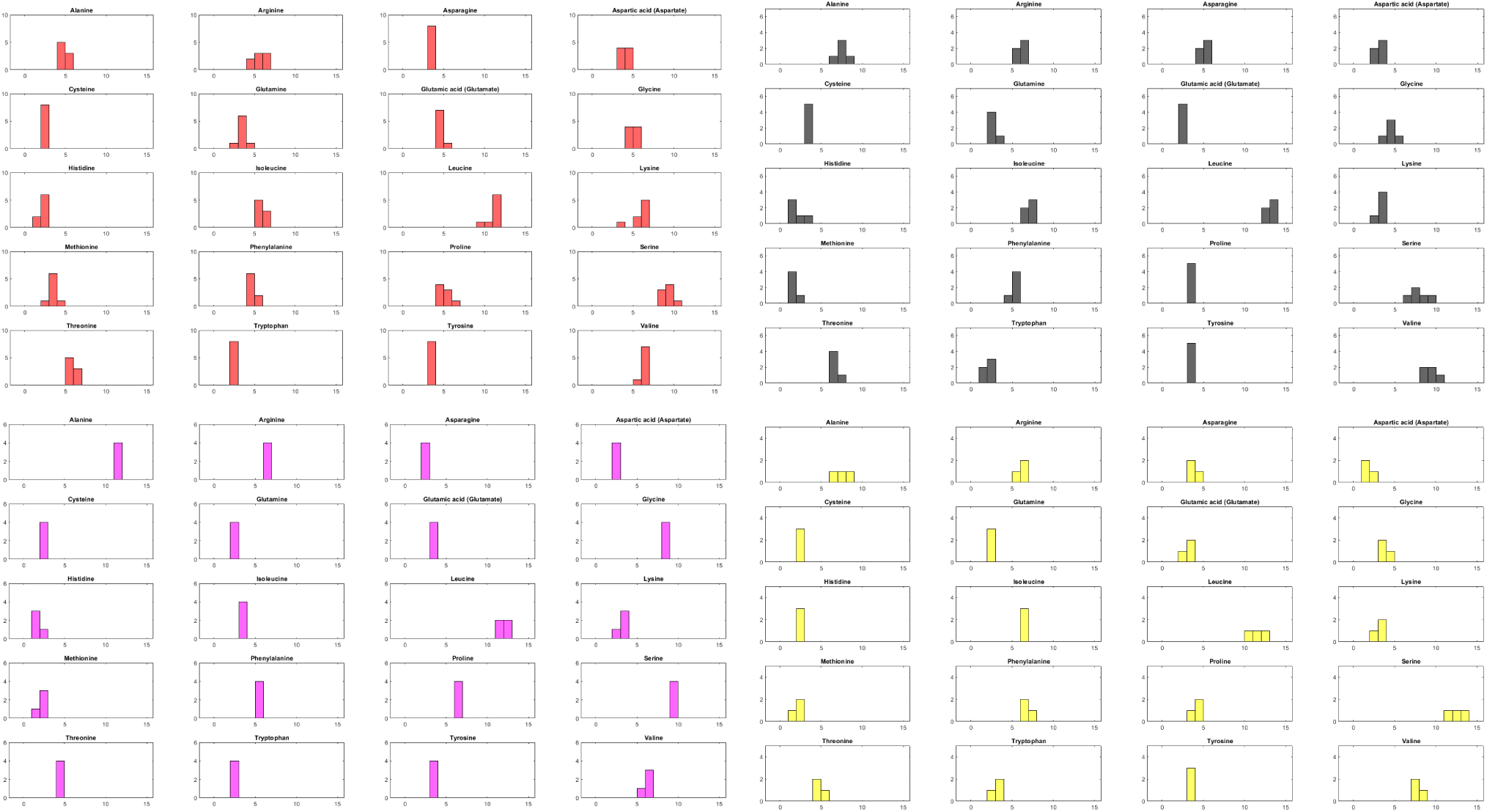
Histograms of amino acid frequency distributions across 20 standard residues. Each subplot represents the count of a specific amino acid, with horizontal axes ranging from 0 to 15 and vertical axes indicating frequency.

**Figure 4:**
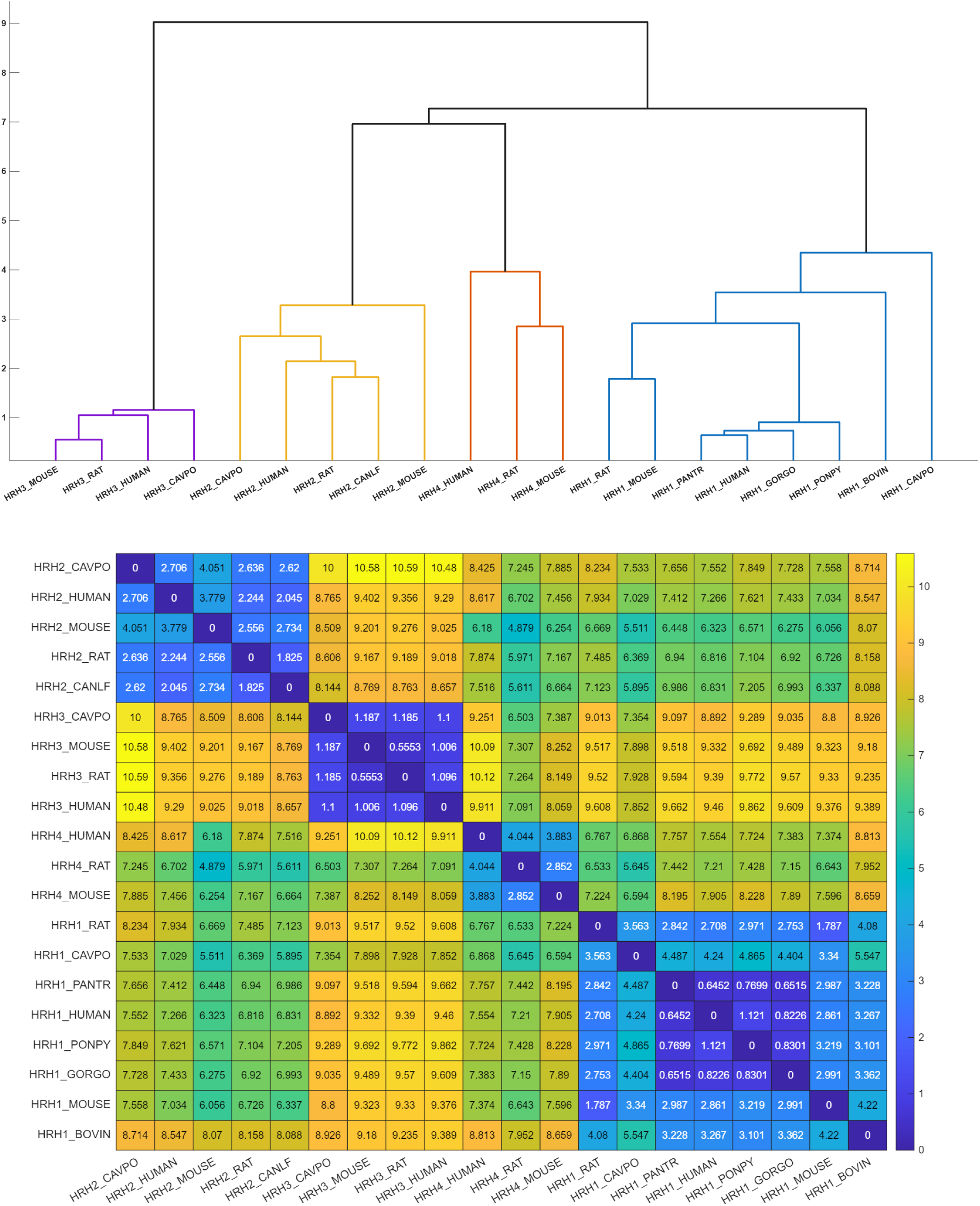
(A) Phylogenetic relationship among all histamine receptors across nine organisms based on amino acid frequency distribution; (B) Heatmap matrix of distance matrix based on relative amino acid frequencies across all histamine receptors.

Although histamine receptors share similarities in activation and binding regions, their protein sequence homology is relatively low, ranging from 16–35%. HRH4 shows moderate conservation among rodents, with rat, mouse, and guinea pig sequences sharing approximately 69%, 68%, and 65% homology, respectively, relative to the human HRH4 sequence [20, 21, 22, 23, 24, 25].

In this study, we applied a suite of quantitative approaches—including sequence homology assessment, identification of conserved residues, amino acid frequency profiling, Shannon entropy, homogeneous poly-string pattern analysis, and intrinsic protein disorder analysis—to investigate the molecular architecture of the four histamine receptor subtypes, belonging to GPCR class A. Although the four HRH1–HRH4 receptors share comparable lengths and overall composition, our analyses highlight subtle yet structurally significant distinctions that define each subtype’s biological role. While individual receptors have been examined qualitatively, a comprehensive quantitative comparison across all four subtypes and multiple mammalian species covering primates, rodents, artiodactyla, and carnivora has been lacking. By integrating evolutionary, structural, and physicochemical metrics into a unified framework, this work addresses that gap. Our multidimensional analysis of conserved versus variable regions, disorder tendencies, compositional signatures, and physicochemical relationships provides a coherent view of evolutionary pressures and subtype-specific features. Overall, this study delivers the first broad, sequence-based quantitative characterization of all four histamine receptors, offering insights directly relevant to the rational design of next-generation antihistamines.

## 2. Data acquisition

Amino acid sequences of Histamine H1 receptor (HRH1), Histamine H2 receptor (HRH2), Histamine H3 receptor (HRH3), and Histamine H4 receptor (HRH4) were retrieved from the UniProtKB using curated accession identifiers for each receptor and organisms (Table 1) ([26]). The dataset comprised a total of 23 sequences distributed across four histamine receptor types: HRH1 (8 sequences), HRH2 (8 sequences), HRH3 (4 sequences), and HRH4 (3 sequences) from nine organisms (Table 2) ([27]). All sequences were downloaded from uniProt in FASTA format and verified for completeness and annotation consistency [27].

**Table 1:**
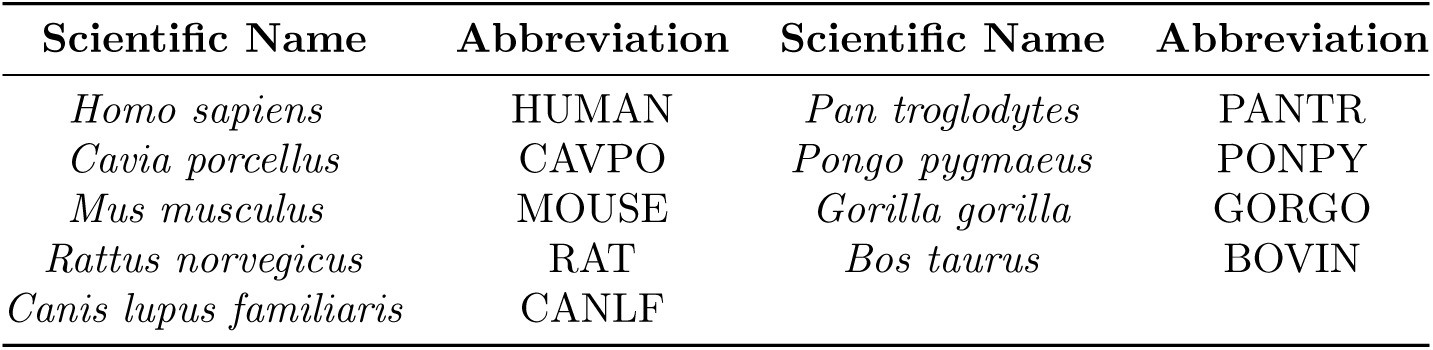
Organism abbreviations and corresponding scientific names used in histamine receptors.

**Table 2:**
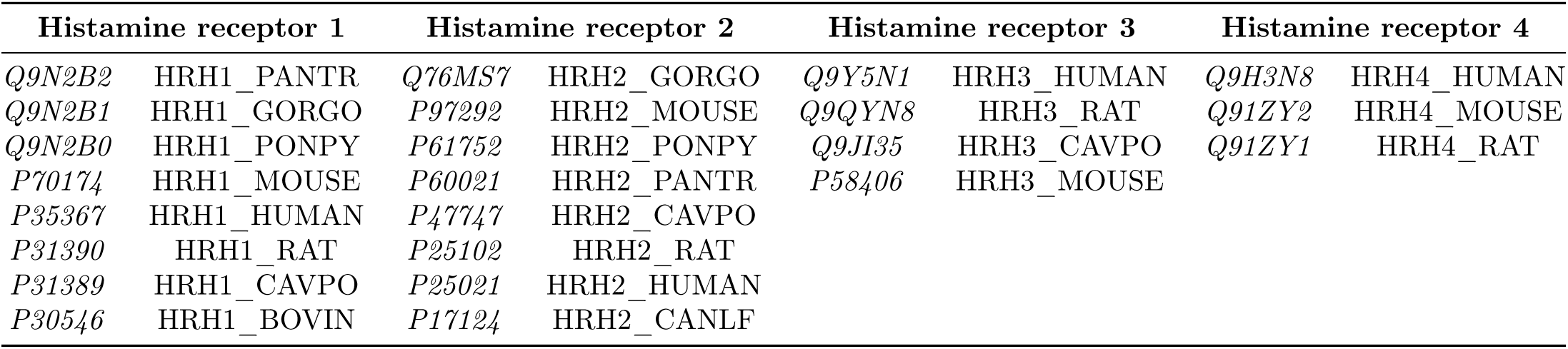
List of histamine receptors across nine organisms.

## 3. Methods

### 3.1. Amino acid sequence homology and phylogeny

Amino acid sequence homology among histamine receptor sequences was assessed using multiple sequence alignment performed with Clustal Omega [28, 29, 30, 31]. This analysis enabled the identification of invariant residues and amino acid residue variations [32].

### 3.2. Determining amino acid frequency composition in histamine receptors

The amino acid frequency, defined as the number of occurrences of each amino acid within a protein sequence, was calculated for all histamine receptor sequences analyzed in this study [33, 34, 35, 36]. To account for differences in sequence length, the relative frequency of each amino acid was obtained by dividing its absolute count by the length of the corresponding receptor sequence and multiplying by 100. This measure reflects the percentage composition of each amino acid within the sequence. Consequently, each histamine receptor sequence is represented as a 20-dimensional vector, corresponding to the relative frequencies of the 20 standard amino acids.

#### 3.2.1. Shannon entropy of histamine receptors based on amino acid frequency distribution

To quantify compositional diversity in histamine receptor sequences, we computed sequence-level Shannon entropy from amino acid frequency distributions. Let *p_i_* denote the relative frequency of amino acid type *i* ∈ {1*, . . .,* 20}; the Shannon entropy *H* of the sequence is then defined as:

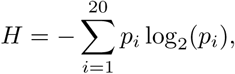

with the convention that terms with *p_i_* = 0 contribute 0 to the sum. Entropy ranged from 0 (complete compositional conservation) to a theoretical maximum of 4.322 for a uniform distribution over 20 amino acids [37, 38, 39, 40].

### 3.3. Determining homogeneous poly-string frequency of amino acids in histamine receptor sequences

A homogeneous poly-string of length *n* is defined as a subsequence composed of *n* consecutive repetitions of the same amino acid [41, 42]. For example, in the sequence “NNNWWNWW”, there exists one homogeneous poly-string of ‘N’ with length 3, another of ‘N’ with length 1, and two homogeneous poly-strings of ‘W’, each with length 2. In identifying homogeneous poly-strings of a given length *n*, only those with exactly length *n* are considered; longer strings are not partitioned into smaller units. To systematically examine these poly-strings across all histamine receptor sequences, we first determined the maximum length of homogeneous poly-strings observed for any amino acid within the dataset. Using this maximum as a reference, we then enumerated the number of homogeneous poly-strings for each amino acid at every possible length from 1 up to the maximum in each protein sequence [42].

### 3.4. Evaluating intrinsic protein disorder in histamine receptor sequences

The intrinsic disorder propensity of histamine receptor sequences was assessed using a suite of well-established per-residue disorder predictors available through the Rapid Intrinsic Disorder Analysis Online (RIDAO) platform. Among these, results from the VSL2B predictor were selected for subsequent analyses [43, 44, 45].

Disorder scores for each residue range from 0 (completely ordered) to 1 (completely disordered). Residues with scores greater than 0.5 are classified as disordered (‘D’). Scores between 0.25 and 0.5 correspond to highly flexible residues (‘HF’), while scores from 0.1 to 0.25 indicate moderately flexible residues (‘MF’). Residues with scores below 0.1 are grouped into the ‘other’ (‘O’) category [43, 46].

### 3.5. Phylogenetic analysis of histamine receptor sequences based on physicochemical profiling

Determining the physicochemical properties of proteins provides important insights into their structural organization, biological functions, stability, and molecular interactions. In the present study, the structural features of histamine receptor sequences were evaluated using the Multiple protein profiler 1.0 (MPP), an integrated tool developed for systematic profiling of protein properties [47]. For each histamine receptor sequence, MPP computed the following properties: molecular weight, theoretical isoelectric point (pI), aliphatic index, instability index, grand average of hydropathicity (GRAVY), aromaticity, and charge distribution. This integrated profiling approach provided a comprehensive overview of the structural features of histamine receptors, enabling downstream interpretation of their functional diversity and stability.

### 3.6. Formation of distance matrices and dendrograms

Multiple sequence alignment based distance matrix was obtained by subtracting similarity matrix from 100. Euclidean distances were computed between all possible pairs of histamine receptor sequences based on relative frequency of amino acids (dimension 20) and features obtained from MPP (dimension 7). Each of the MPP based feature was normalized and both global phylogenies (encompassing all histamine receptor sequences) as well as individual phylogenies for each receptor subtype (H1, H2, H3, and H4) were constructed. This dual-level approach enabled the identification of inter and intra subtype-specific clustering patterns across histamine receptors based on physicochemical properties.

## 4. Results and analyses

### 4.1. Amino acid sequence homology of histamine receptors across mammals

In determining sequence homology across histamine receptors of all four subtypes in nine organisms, hierarchical clustering was performed on the full-length amino acid sequences of HRH1, HRH2, HRH3, and HRH4 (Table 1) [35, 48, 42, 49, 41].

Complete sequence identity (100%) was detected among HRH2 sequences from *Homo sapiens*, *Pan troglodytes*, *Pongo pygmaeus*, and *Gorilla gorilla* (HRH2_HUMAN, HRH2_PANTR, HRH2_PONPY, and HRH2_GORGO), highlighting strong evolutionary conservation within primates (Figure 5). HRH2 sequences of RAT, MOUSE and CAVPO, all belong to Rodent order, were not identical. All pairwise differences among HRH1 sequences of the four primate organisms (HUMAN, PANTR, PONPY, GORGO) ranged from 97% to 99%. Based on eight HRH1 sequences, HUMAN was closest to PANTR and farthest from CAVPO with 98.77% and 73.43% similarities respectively. Similarity between RAT and MOUSE was highest for HRH3 (98.88%) and lowest for HRH4 (85.17%). Among all four subtypes of receptors, similarity between HUMAN and both RAT and MOUSE was maximum (around 93%) and minimum (approximately 68%) based on HRH3 and HRH4 respectively. Thus, HRH4 sequences demonstrated notably more divergence among different organisms than other receptors. Comparing histamine receptors of subtypes 1, 2 and 3, similarity between CAVPO and RAT as well as CAVPO and MOUSE was highest (around 95%) based on HRH3 and lowest (around 71%) based on HRH1. CAVPO was around 2% closer to HUMAN than RAT or MOUSE based on HRH1 and HRH2 while 2% more distant based on HRH3. Among six possible pairs of HUMAN histamine receptors, maximum similarity was found between HUMAN_HRH3 and HUMAN_HRH4 (40.11%) followed by HUMAN_HRH1 and HUMAN_HRH2 (32.78%). Other pairs had values between 23% and 32.78%. Similar observations were found for MOUSE and RAT.

**Figure 5:**
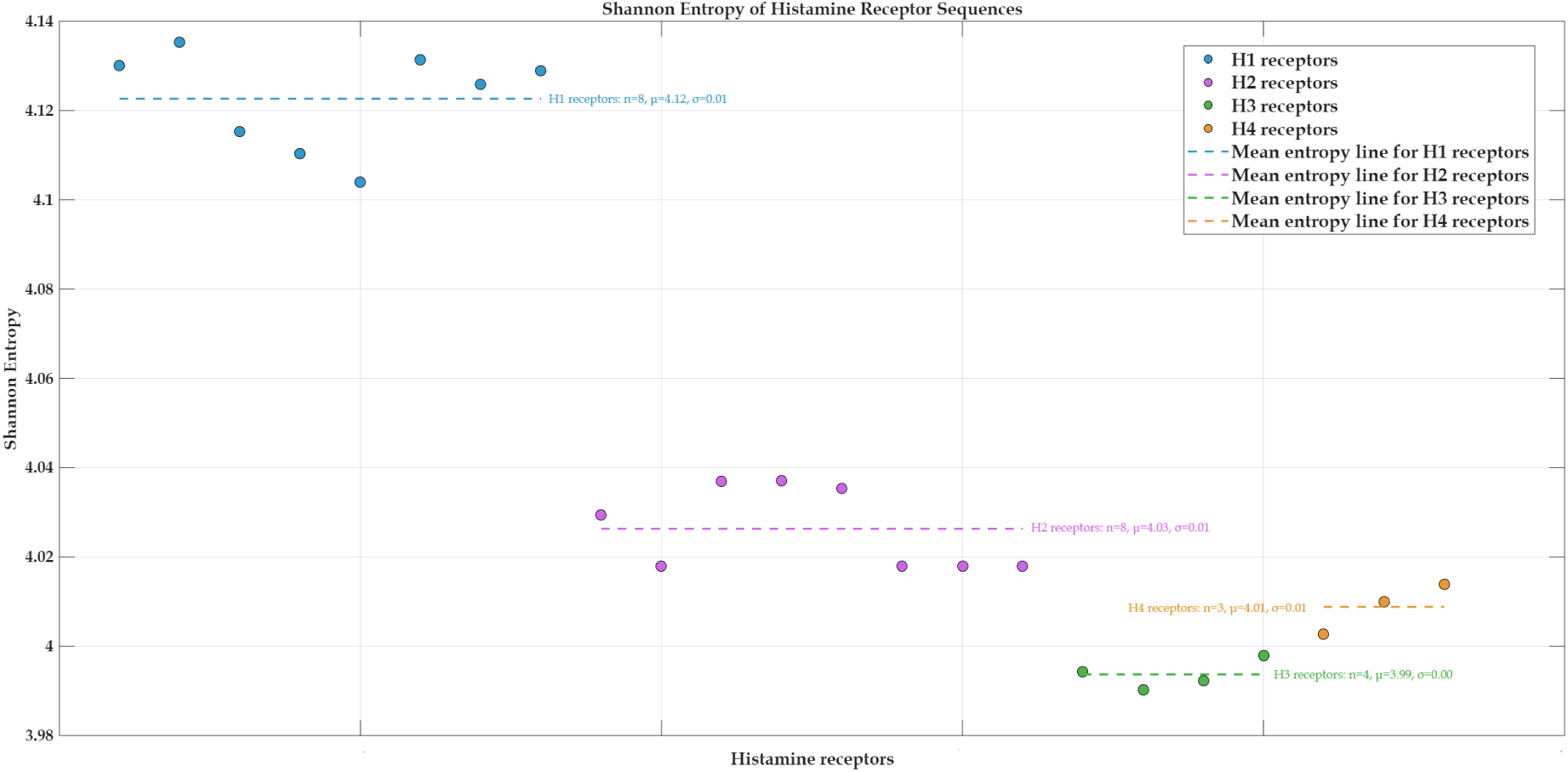
Scatter plot of Shannon entropy values for individual histamine receptor sequences (H1–H4). Each receptor subtype is color-coded, and dashed horizontal lines indicate group-wise mean entropy. The plot highlights subtype-specific differences in sequence complexity, with H1 showing the highest entropy and H4 the lowest.

#### 4.1.1. Amino acid invariance and amino acid residue changes across all histamine receptors

Multiple-sequence alignment of all mammalian histamine receptors—including H1, H2, H3, and H4 receptor sequences from human, mouse, rat, rabbit, bovine, canine, guinea pig, and non-human primates—revealed a distinct set of invariant amino acid residues that remained conserved across both species and receptor subtypes. When these conserved positions were mapped to the human H1R reference sequence (P35367), the invariant residues identified were: N45, V48, A51, L58, L69, D73, G77, P82, W93, C100, D107, S114, D124, R125, V129, Y135, W152, L163, C180, F199, P202, I213, L417, F424, C427, W428, P430, Y431, W455, L456, N460, S461, N464, P465, Y468, and F475 (Table 3). These residues remained unchanged in every mammalian H1–H4 receptor sequence analysed, indicating that they represent deeply conserved structural constraints essential for receptor function [50, 51]. Most invariant residues fell within transmembrane helices (TM1–TM7) and mapped onto regions corresponding to the ligand-binding cavity, the hydrophobic transmembrane core, and universal activation motifs characteristic of Class A GPCRs. All functionally critical residues identified in structural and mutagenesis studies were also conserved across all receptors. These included D107 (TM3), which is indispensable for agonist binding and activation, and W428 and Y431 (TM6), which form part of the classical GPCR micro switch network; whose mutations at those positions abolish histamine-induced activation or induce constitutive activity [52, 53]. The highly conserved activation motif – the NPxxY in TM7 (represented by Y468, N464, and P465) was also invariant across all sequences in a manner consistent with its universal role in receptor activation and stabilization of conformational active state [54]. The strict conservation of N460 and S461 upstream of the NPxxY motif reflects their essential role in the TM7 conformational network that stabilizes the active state of Class A GPCRs [52]. The strong conservation of these positions across mammalian species and histamine receptor subtypes is indicative of great evolutionary pressure to maintain structural integrity, ligand binding specificity, and activation dynamics, as well as G-protein coupling productiveness. Since histamine is an identical endogenous ligand in all mammals, ligand recognition and binding residues like D107, C427, W428, and Y431 cannot tolerate substitutions without compromising receptor function. The invariance of hydrophobic residues L69, W152, L163, L417, W455, and L456 further argues for their role in establishing the GPCR fold since the transmembrane helices of Class A GPCRs require precise packing for folding and membrane insertion to occur [50]. Although no structure of the histamine receptor–G-protein complex is yet available, conserved residues at the cytoplasmic ends of TM3 and TM5 are expected to participate in G-protein coupling, in line with general structural principles of Class A GPCR drawn from GPCR–G-protein complex structures [55]. Taken together, the invariant residues reported in this study represent evolutionarily locked positions defining the structural and functional architecture of the histamine receptor family. Their conservation across both receptor types and mammalian genomes highlights the importance of these proteins in ligand binding, receptor stability, activation, and intracellular signalling. It also illustrates the strong evolutionary pressure operating on histamine signalling pathways.

**Table 3:** List of invariant residues and their respective positions with respect to P35367 (HRH1_HUMAN) across all histamine receptors (all subtypes)

| Sl. No. | Amino acid position | Invariant residue | position | Description | Mutagenesis |
| --- | --- | --- | --- | --- | --- |
| 1 | 45 | N | Transmembrane | Helical; Name=1 |  |
| 2 | 48 | V | Transmembrane | Helical; Name=1 |  |
| 3 | 51 | A | Topological domain | Cytoplasmic |  |
| 4 | 58 | L | Topological domain | Cytoplasmic |  |
| 5 | 69 | L | Transmembrane | Helical; Name=2 |  |
| 6 | 73 | D | Transmembrane | Helical; Name=2 |  |
| 7 | 77 | G | Transmembrane | Helical; Name=2 |  |
| 8 | 82 | P | Transmembrane | Helical; Name=2 |  |
| 9 | 93 | W | Topological domain | Extracellular |  |
| 10 | 100 | C | Transmembrane | Helical; Name=3 |  |
| 11 | 107 | D | Transmembrane | Helical; Name=3 | Change from D to A or E abolishes histamine-mediated activation, indicating that D107 is essential for agonist binding and stabilization of the active receptor conformation [56]. |
| 12 | 114 | S | Transmembrane | Helical; Name=3 |  |
| 13 | 124 | D | Topological domain | Cytoplasmic |  |
| 14 | 125 | R | Topological domain | Cytoplasmic |  |
| 15 | 129 | V | Topological domain | Cytoplasmic |  |
| 16 | 135 | Y | Topological domain | Cytoplasmic | A change from Y to A at position 135 has no effect on histamine-mediated activation, indicating that Y135 is not essential for agonist binding or stabilization of the active receptor conformation [56]. |
| 17 | 152 | W | Transmembrane | Helical; Name=4 |  |
| 18 | 163 | L | Transmembrane | Helical; Name=4 |  |
| 19 | 180 | C | Topological domain | Extracellular |  |
| 20 | 199 | F | Transmembrane | Helical; Name=5 |  |
| 21 | 202 | P | Transmembrane | Helical; Name=5 |  |
| 22 | 213 | I | Topological domain | Cytoplasmic |  |
| 23 | 417 | L | Transmembrane | Helical; Name=6 |  |
| 24 | 424 | F | Transmembrane | Helical; Name=6 | Important for agonist binding |
| 25 | 427 | C | Transmembrane | Helical; Name=6 | Important for agonist binding |
| 26 | 428 | W | Transmembrane | Helical; Name=6 | A change from W to A results in a loss of histamine-mediated activation, indicating that this residue is critical for agonist binding [56]. |
| 27 | 430 | P | Transmembrane | Helical; Name=6 |  |
| 28 | 431 | Y | Transmembrane | Helical; Name=6 | 1. A change from Y to F results in a loss of histamine-mediated activation [56].<br>2. A change from Y to R leads to increased histamine receptor activity, indicating a constitutively active receptor [56]. |
| 29 | 455 | W | Transmembrane | Helical; Name=7 |  |
| 30 | 456 | L | Transmembrane | Helical; Name=7 |  |
| 31 | 460 | N | Transmembrane | Helical; Name=7 |  |
| 32 | 461 | S | Transmembrane | Helical; Name=7 |  |
| 33 | 464 | N | Transmembrane | Helical; Name=7 |  |
| 34 | 465 | P | Transmembrane | Helical; Name=7 |  |
| 35 | 468 | Y | Transmembrane | Helical; Name=7 |  |
| 36 | 475 | F | Topological domain | Cytoplasmic |  |

An analysis of amino acid changes in all mammalian histamine receptors from H1 to H4, based on the human H1 receptor sequence P35367, showed a systematic pattern of divergence in comparison to the highly conserved invariant positions identified earlier (Table 4).The changes shown in Table 4 took place only in non-essential, flexible residues, while invariants surveyed the critical elements of Class A GPCRs. HRH2 receptors exhibited a consistent signature of shared substitutions—including L30I/V, V76L, W103Y, S128A, L154I, F156I, Y214F, A414T and F479Y (among itself)—including E55N, I75L, L95F, V109M, V118L, I120M, L121I, I148L etc (along with H3 and H4)—reflecting the subtype’s divergence toward Gs-coupling and cAMP signaling, distinct from the Gq/11-driven H1R or the Gi/o-linked H3/H4 receptors. In contrast, HRH3 and HRH4 shared extensive parallel changes such as L34M, I66L/F, V67L, L74F, L133V, S155A, M206V, Y458W, L463V and N472H highlighting their closer evolutionary relationship and functional similarity. The majority of the substitutions corresponds to conservative amino acid substitutions, Leu/Val, Ile/Leu, Phe/Tyr, and so on, suggesting that there is tolerated variability at these sites that does not affect the overall fold of the GPCRs. Alignment of these positions to the H1 receptor crystalline structure indicates that all the variable residues are found in lipid-facing TM positions, the extracellular loops, or the intracellular loops, all of which tend to have a rapid rate of evolution and do not affect the conserved ligand-binding site [50, 51, 52]. The lack of substitutions in functionally essential positions such as D107, C427, W428, and Y431 further reinforces the idea that the activation process of histamine receptors remains extremely well-conserved across mammals. In sum, the data provided in Table 4 show the degree of amino acid variation that has occurred due to the use of permissive sites for the diversification of histamine receptors across the tree of life, with the structural framework that remains critically well-conserved.

**Table 4:**
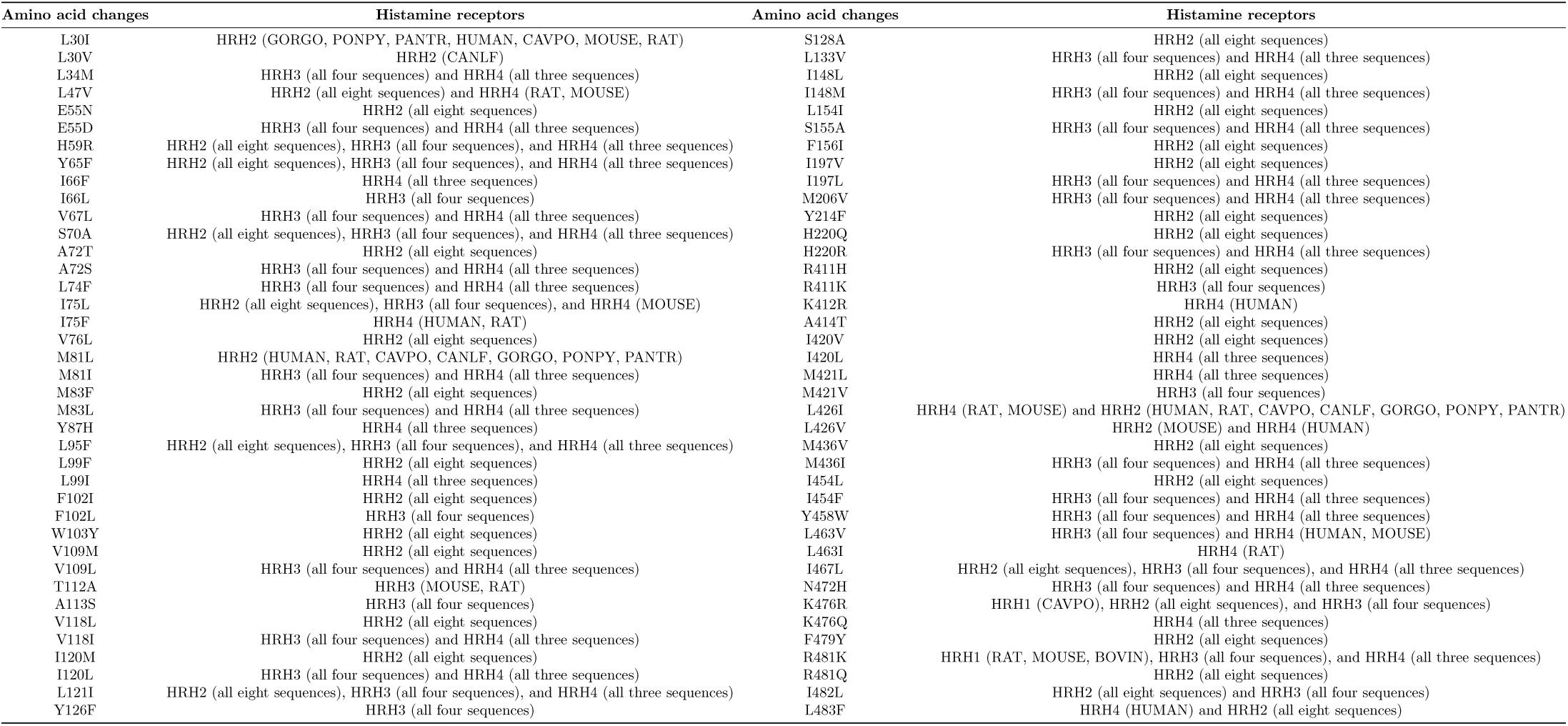
List of amino acid changes (*strong conservation*—amino acids at that position are *highly similar* in physicochemical properties, such as charge, polarity, or hydrophobicity) across all histamine receptors (all subtypes) with reference to P35367 (HRH1_HUMAN)

4.2. *Amino acid frequency distribution and associated phylogenetic relationship among histamine receptors*

#### 4.2.1. Amino acid frequency distribution among histamine receptors

Across HRH1, HRH2, and HRH3 sequences, Leucine consistently exhibited the highest median percentages (11.29%, 13.09%, and 12.02%, respectively). In HRH1, Serine ranked second (median 9.2%), while all other amino acids had values below 6.98%. In HRH2, Valine followed Leucine (median 9.07%), except in HRH2_MOUSE where Serine exceeded Valine by 0.25%. HRH1 and HRH2 contained similar amounts of Glycine, Phenylalanine, Tryptophan, and Tyrosine.

In HRH3 sequences, Alanine was the second most abundant residue (median 11.12%), markedly higher than in HRH1 (median 4.93%), indicating substantial inter-subtype variation. Asparagine, Cysteine, and Tryptophan levels were identical across all four HRH3 sequences. Median values of Cysteine, Phenylalanine, Tyrosine, and Valine were comparable between HRH1 and HRH3, apart from Serine. Isoleucine and Asparagine showed the lowest medians in HRH3 (3.6% and 2.47%) and the highest in HRH2 (7.15% and 5.2%) among four subtypes. Proline was significantly lower in HRH2 (median 3.33%) compared to HRH3 (median 6.35%). Methionine levels were comparable between HRH2 and HRH3, whereas Tyrosine displayed a different pattern.

HRH4 diverged from the other subtypes. In HRH4_HUMAN and HRH4_MOUSE, Serine was most abundant followed by Leucine, while HRH4_RAT contained 0.5% more Leucine than Serine. Glycine differed markedly between HRH3 (median 8.65%) and HRH4 (median 3.58%). Lysine was elevated in HRH1 (median 6.78%) compared to the other subtypes, except HRH1_CAVPO which contained markedly less Lysine (3.69%). Tyrosine was present in nearly the same amount (∼3.6%) across all sequences of the four subtypes.

Furthermore, across the four HRH subtypes, Alanine exhibited the highest standard deviation among the twenty amino acids in each in each of HUMAN, MOUSE, and RAT.

#### 4.2.2. Phylogenetic reletionship among histamine receptors based on amino acid frequency distributions

Among all possible pairs across the four HRH subtypes, the highest pairwise distance (10.589) was observed between HRH3_RAT and HRH2_CAVPO. Within HRH2 sequences, the greatest distance (4.051) occurred between HRH2_MOUSE and HRH2_CAVPO. Intragroup pairwise distances were lowest for HRH3 compared to the other three subtypes, with the maximum value (1.187) noted between HRH3_MOUSE and HRH3_CAVPO. For HRH1 sequences, the highest distance (5.547) was found between HRH1_BOVIN and HRH1_CAVPO.

In HUMAN sequences, the largest inter-subtype distance (9.91) was detected between HRH3 and HRH4, primarily due to Glycine and Alanine being 5.21% and 4.83% more abundant in HRH3, respectively. The lowest distance (7.266) was observed between HRH1 and HRH2. In MOUSE sequences, the highest distance (9.323) was revealed between HRH3 and HRH1, driven mainly by Alanine and Glycine, which were 6.09% and 4.48% more abundant in HRH3, respectively. The lowest distance (6.056) was observed between HRH1 and HRH2. In RAT sequences, the greatest distance (9.52) was found between HRH3 and HRH1, primarily due to Alanine and Glycine being 6.5% and 4.24% more abundant in HRH3, respectively. The lowest distance (5.971) was observed between HRH4 and HRH2.

#### 4.2.3. Shannon entropy across histamine receptors

Quantifying sequence complexity across histamine receptor of all four subtypes involved calculating Shannon entropy for each protein sequence in the HRH1, HRH2, HRH3, and HRH4 receptors. The resulting Shannon entropy revealed subtype-specific differences in amino acid diversity [41, 42, 49].

H1 receptors showed the highest average entropy (*µ* ≈ 4.123), indicating a greater degree of sequence variability [42]. H2 sequences had 2% lower entropy than H1 (*µ* ≈ 4.033), suggesting a more constrained sequence pattern. H3 and H4 receptors displayed the lowest entropy values (*µ* ≈ 4.008 and *µ* ≈ 4.005, respectively). This downward trend in entropy—H1 *>* H2 *>* H3 ≈ H4—reflects a gradual reduction in sequence complexity across receptor subtypes.

### 4.3. Polystring frequency distribution across histamine receptors of all subtypes

Polystring analysis across histamine receptor subtypes revealed distinct patterns of amino acid repetition [35, 41]. For histamine receptor subtype 1 (HRH1), the tryptophan polystring of length three (‘WWW’) was consistently observed in all eight sequences with a count of 1, except for HRH1_BOVIN, which exhibited a count of 2 (Table 5). Among these eight HRH1 sequences, the alanine polystring ‘AAA‘ was uniquely present in HRH1_HUMAN (**Supplementary file 1**). Notably, HRH1_HUMAN also exhibited the highest frequency of polystring occurrence of length 3, with a total count of four. Arginine polystrings of length 2 (RR) were uniformly present across all eight HRH1 sequences with a count of 1. In contrast, polystrings of Leucine, Lysine, Tyrosine, and Valine of length 2 were absent in all HRH1 sequences (**Supplementary file 1**).

**Table 5:** Frequency of homogeneous poly-string in histamine receptor sequences.

| H1 receptors | l=1 | l=2 | l=3 |
| --- | --- | --- | --- |
| HRH1_RAT | 437 | 23 | 1 |
| HRH1_CAVPO | 432 | 25 | 2 |
| HRH1_PANTR | 435 | 21 | 3 |
| HRH1_HUMAN | 435 | 20 | 4 |
| HRH1_PONPY | 438 | 20 | 3 |
| HRH1_GORGO | 436 | 21 | 3 |
| HRH1_MOUSE | 445 | 20 | 1 |
| HRH1_BOVIN | 444 | 19 | 3 |

| H3 receptors | l=1 | l=2 | l=3 | l=4 | l=5 | l=7 |
| --- | --- | --- | --- | --- | --- | --- |
| HRH3_CAVPO | 355 | 35 | 5 | 0 | 1 | 0 |
| HRH3_MOUSE | 356 | 32 | 7 | 1 | 0 | 0 |
| HRH3_RAT | 354 | 33 | 7 | 1 | 0 | 0 |
| HRH3_HUMAN | 354 | 33 | 3 | 1 | 1 | 1 |

| H2 receptors | l=1 | l=2 | l=3 |
| --- | --- | --- | --- |
| HRH2_CAVPO | 327 | 13 | 2 |
| HRH2_HUMAN | 319 | 17 | 2 |
| HRH2_MOUSE | 345 | 23 | 2 |
| HRH2_RAT | 314 | 19 | 2 |
| HRH2_CANLF | 319 | 17 | 2 |

| H4 receptors | l=1 | l=2 | l=3 |
| --- | --- | --- | --- |
| HRH4_HUMAN | 348 | 21 | 0 |
| HRH4_RAT | 353 | 19 | 0 |
| HRH4_MOUSE | 348 | 20 | 1 |

All five sequences corresponding to histamine receptor subtype 2 (HRH2) contained one instance each of the polystrings ‘III’ and ‘WWW’. Polystrings of Cysteine, Histidine, Leucine, and Tyrosine of length 2 were absent across all five HRH2 sequences (**Supplementary file 1**). A single occurrence of the polystring ‘NN‘ was consistently noted in each of the five HRH2 sequences (Table 5).

Within histamine receptor subtype 3 (HRH3), a polystring of length seven was observed exclusively in the human sequence, corresponding to the amino acid Glutamine. No polystrings of length six were detected in any of the four organisms. HRH3_CAVPO and HRH3_HUMAN exhibited one instance each of ‘AAAAA’ and ‘MMMMM’, respectively. Alanine was the amino acid associated with polystrings of length four. The polystring ‘SSS’ was present in all four HRH3 sequences with a count of 2. Additionally, the polystrings ‘CC’, ‘KK’, ‘FF’, ‘PP’, and ‘WW’ were consistently observed across all four sequences with counts of four, one, one, three, and one, respectively. Single-length polystrings of Arginine (‘R’), Lysine (‘K’), and Tyrosine (‘Y’) were found with counts of twelve, nine, and thirteen, respectively, in all four organisms. Importantly, no polystrings of Arginine, Asparagine, Aspartic acid, Leucine, Threonine, Tyrosine, or Valine with lengths greater than one were present in any of the HRH3 sequences (Table 5).

For histamine receptor subtype 4 (HRH4), a polystring of length three corresponding to ‘SSS’ was observed in HRH4_MOUSE. No polystrings of length two corresponding to Arginine, Asparagine, Glutamine, Glutamic acid, Leucine, Lysine, Phenylalanine, Tyrosine, or Valine were present in any of the three HRH4 sequences. The polystring ‘PP’ was consistently observed with a count of three across all three HRH4 sequences (Table 5).

### 4.4. Intrinsic protein disorder among histamine receptors

G protein-coupled receptors (GPCRs) contain significant levels of intrinsic disorder [57]. This is expected, as the human GPCR-G protein signaling system—comprising over 800 diverse GPCRs and numerous G proteins in humans—must recognize a wide range of extracellular ligands and initiate a variety of intracellular signaling cascades in response. This structural disorder facilitates the multifunctionality and binding promiscuity of these receptors, allowing a single GPCR to recognize multiple extracellular signals and interact with more than one G protein, enabling different receptors to converge on the same G protein or permitting one ligand to activate several different GPCRs. As illustrated by Figure 6, human histamine receptors demonstrate this characteristic, with high levels of predicted disorder being primarily localized in their intracellular C-terminal tails.

**Figure 6:**
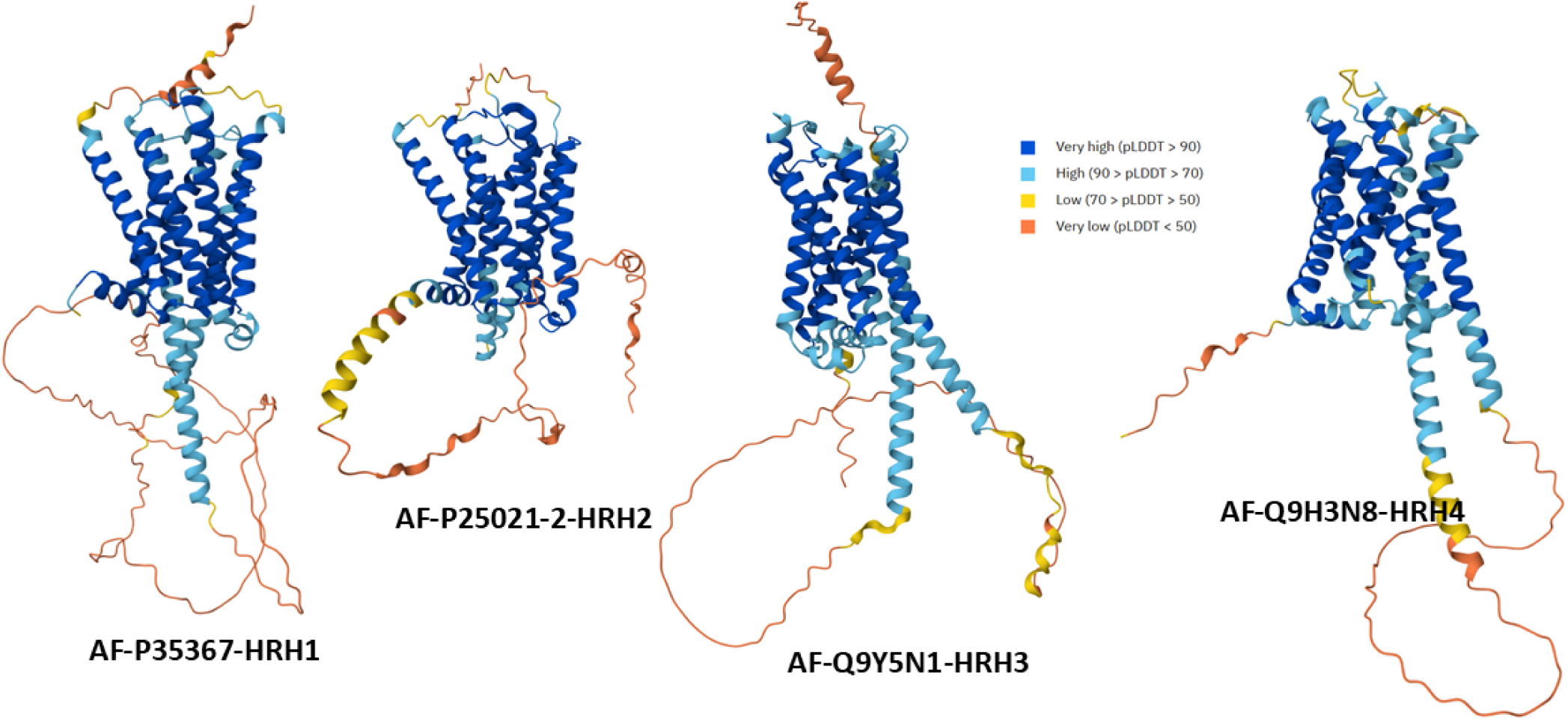
3D structures modeled for human histamine receptors HRH1, HRH2, HRH3, and HRH4 by AlphaFold [58]. The structure is colored based on the per-residue model confidence (pLDDT), which ranges between 0 and 100. Regions with very high (pLDDT > 90), high (90 > pLDDT > 70), low (70 > pLDDT > 50), and very low model confidence (pLDDT < 50) are shown by blue, cyan, yellow, and orange colors, respectively. Regions with low pLDDT may be unstructured in isolation.

**Figure 7:**
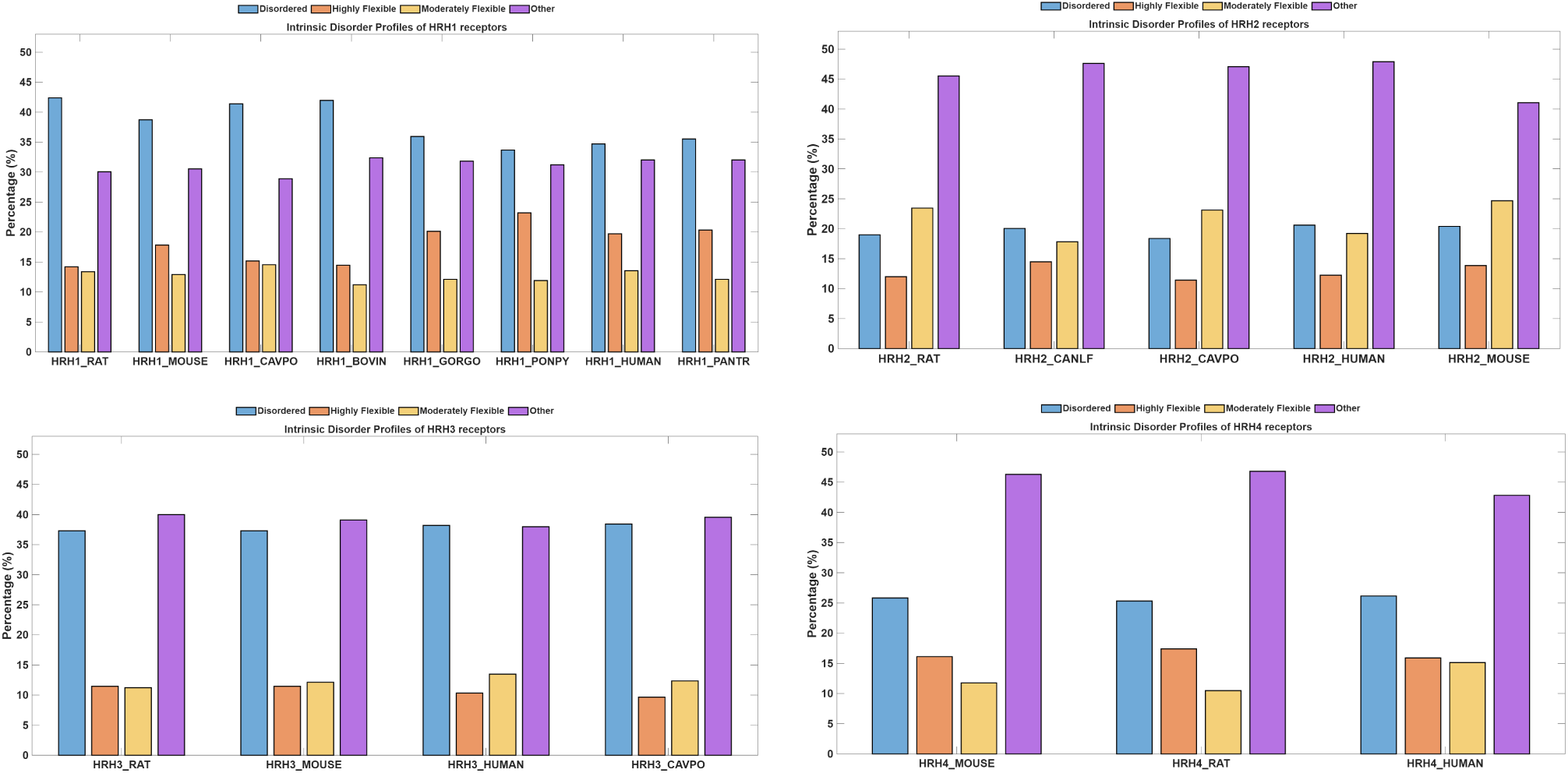
Percentages of disordered, highly flexible, moderately flexible and other residues based on respective Intrinsic protein disorder profiles.

The ratio of the percentage of disordered residues to other residues was highest in HRH1 (with mean 1.226), followed by HRH3 (with mean 0.966), and lowest in HRH2 (with mean 0.431). Similarly, the ratio of highly flexible to moderately flexible residues was greatest in HRH1 (with mean 1.44), followed by HRH4 (with mean 1.36), and lowest in HRH2 (with mean 0.603). The maximum percentage of highly flexible residues was observed in HRH1 (with mean 18.13%), whereas the minimum was obtained in HRH3 (with mean 10.73%). The highest percentage of other residues was observed in HRH4 receptors.

### 4.5. Phylogenetic relationship among histamine receptors based on physiochemical properties

Phylogenetic clustering based on pairwise physicochemical distances revealed distinct subtype-specific cohesion among histamine receptors (Figure 8) [35]. HRH3 sequences formed the tightest cluster, with HRH3_MOUSE and HRH3_CAVPO exhibiting the highest similarity, followed closely by HRH3_HUMAN and HRH3_RAT, indicating strong conservation of biophysical properties within this HRH3 subtype. HRH1 receptors displayed broader intra-subtype dispersion, yet maintained cohesive subgroupings: HRH1_HUMAN clustered closely with HRH1_PONPY and HRH1_GORGO, HRH1_BOVIN with HRH1_PANTR, and HRH1_RAT with HRH1_MOUSE (Figure 8). These pairings suggest lineage-specific conservation of structural features, particularly among primates and rodents. HRH2 sequences, while internally cohesive-evidenced by close pairing of HRH2_HUMAN with HRH2_RAT and HRH2_CANLF, exhibited the greatest overall separation from other subtypes, consistent with their distinct gastric signaling roles and divergent physicochemical profiles (Figure 8). Collectively, the dendrogram topology supports subtype-specific clustering driven by conserved biophysical constraints, with HRH3 showing the highest internal similarity and HRH2 forming the most isolated clade.

**Figure 8:**
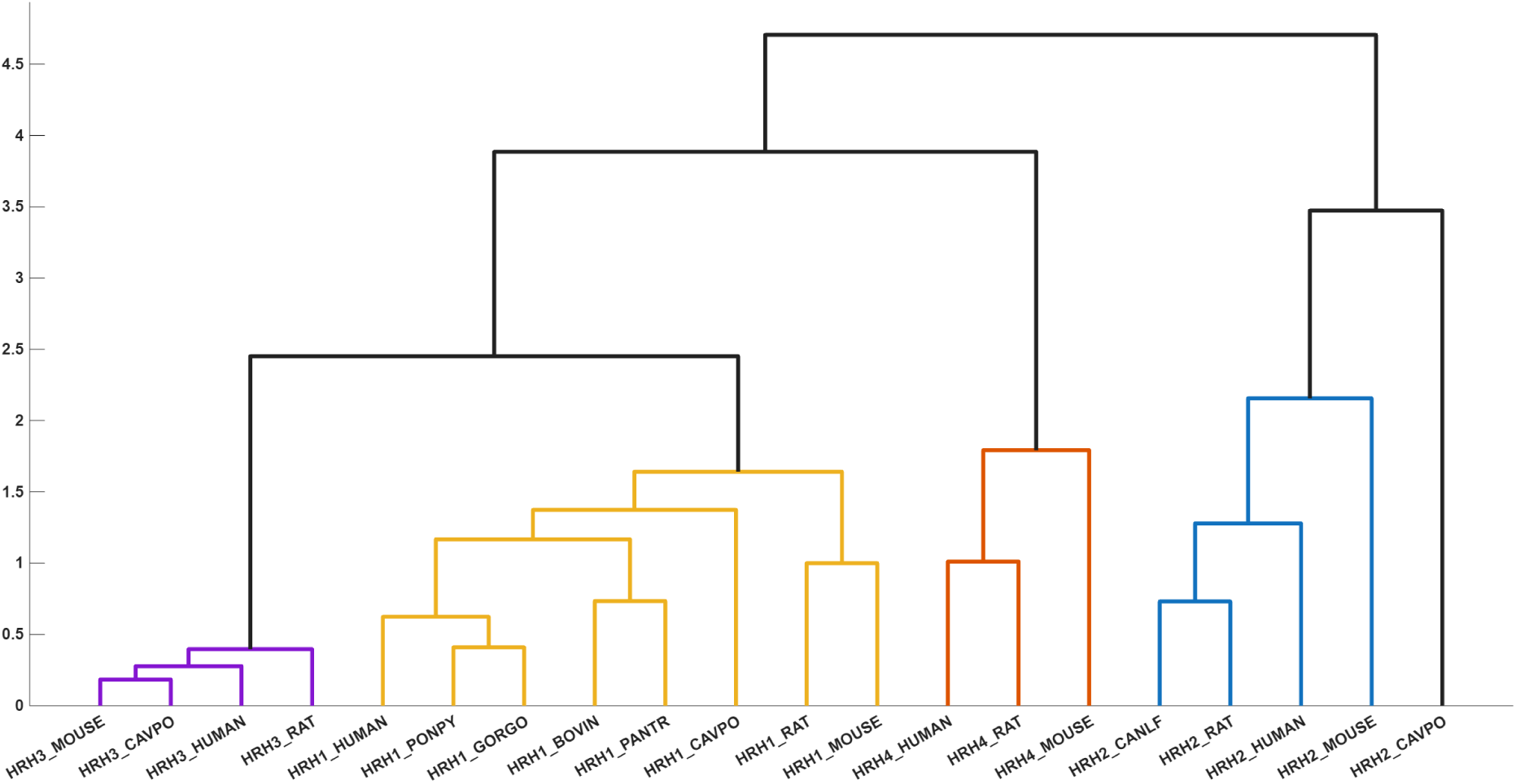
Phylogeny of histamine receptors derived from the physiochemical properties

Comprehensive phylogenetic analysis of histamine receptor subtypes (HRH1–HRH4) across multiple mammalian species revealed distinct patterns of intra-subtype cohesion and interspecies divergence. HRH1 receptor sequences exhibited lineage-specific clustering, with higher primates (HRH1_HUMAN, HRH1_PONPY, HRH1_PANTR, HRH1_GORGO) forming a tightly linked clade, indicative of strong conservation in receptor architecture (Figure 9). HRH1_BOVIN clustered proximally with HRH1_GORGO, suggesting partial structural similarity between bovine and primate lineages. Rodent sequences (HRH1_RAT, HRH1_MOUSE) formed a distinct subclade, joined by HRH1_CAVPO at a slightly higher branch point, reflecting moderate divergence (Figure 9). Phylogenetic analysis of HRH2 receptor sequences across five mammalian species revealed subtype-specific conservation with internal subgroupings. HRH2_CANLF and HRH2_RAT formed the closest pair, with HRH2_HUMAN joining this cluster at a slightly higher branch point. HRH2_MOUSE and HRH2_CAVPO formed a second compact pair (Figure 9). The overall topology supports a cohesive HRH2 clade with two tight subgroups and a modest separation between them. HRH3 receptors revealed a compact topology, with HRH3_MOUSE and HRH3_HUMAN forming the closest pair, followed by HRH3_RAT and the more distantly related HRH3_CAVPO, suggesting minimal intra-subtype dispersion and species-specific gradation (Figure 9). HRH4 receptors formed a cohesive clade, with HRH4_HUMAN and HRH4_RAT exhibiting the highest similarity, and HRH4_MOUSE joining at a slightly elevated branch point, consistent with conserved immune signaling roles (Figure 9). Collectively, the dendrogram topologies underscore subtype-specific conservation, with HRH3 and HRH4 showing the tightest clustering, HRH1 displaying lineage-dependent dispersion, and HRH2 forming the most structurally distinct clade.

**Figure 9:**
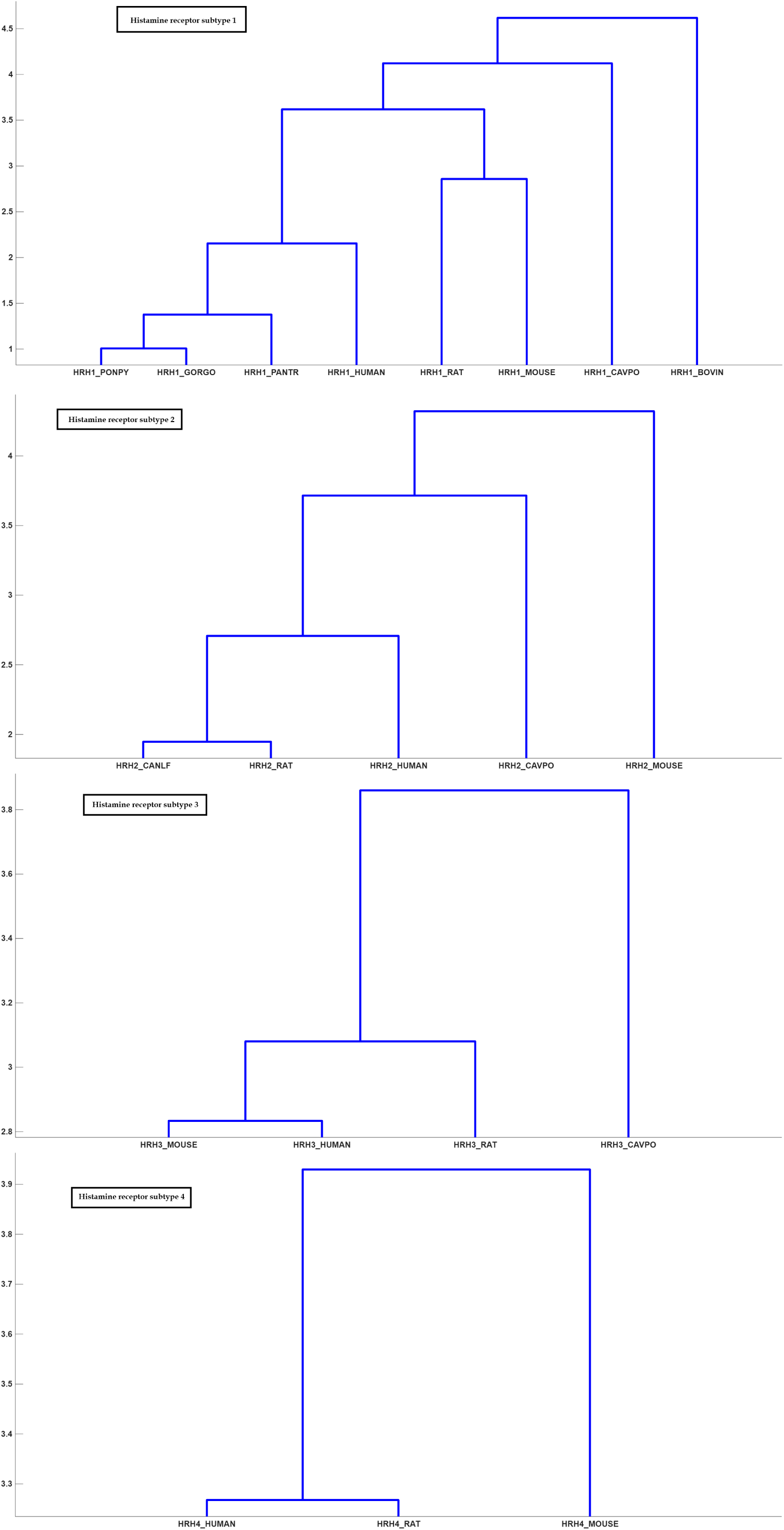
Phylogeny of individual set of histamine receptors derived from the physiochemical properties

## 5. Discussion

This study provides a comprehensive comparative analysis of mammalian histamine receptors (HRH1–HRH4), integrating sequence homology, invariant residues, amino acid substitutions, compositional frequencies, entropy, polystring distributions, intrinsic disorder, and phylogenetic clustering. Together, these results highlight a dual evolutionary strategy: strict conservation of structural cores essential for ligand recognition and activation, alongside variability at permissive positions that enable subtype specialization and species-level adaptation.

Homology analysis reveals that there is significant evolutionary pattern variation among the histamine receptors of mammalian species. HRH2 is the most conserved type of histamine receptors, and the sequences of primates are perfectly identical. This is due to the high functional constraints of this receptor type on gastric and neural transmissions [8]. HRH1 and HRH3 are also conserved to a high degree among primates and rodents, showing the high functional relevance of these two receptor types. HRH4 is the most divergent among the four types of histamine receptors. This indicates that this receptor type has evolved at a faster rate because of the species-specific variation of the immune response among different species [24].

The multiple sequence alignment has identified a short but highly conserved core of invariant amino acids common to all four histamine receptor subtypes and all mammalian species analysed. These residues are primarily centered upon the transmembrane helices and represent core functional elements of Class A GPCRs, such as the ligand-binding pocket, the hydrophobic core, and the universal activation helices, for instance, the NPxxY motif [50, 51, 52, 53, 59]. The full conservation of these amino acids, particularly those that are critical to functionality, as present in D107, W428, and the Y431 position, suggests intense evolutionary pressure to retain the basic mechanisms involved in histamine recognition and activation and G-protein coupling [50]. In contrast, the amino acid substitutions that were identified across species were restricted to positions that are either non-essential or flexible. The substitutions that were identified were largely conservative, occurring at lipid-exposed surfaces of the transmembrane domain, or at extracellular or intracellular loops, the position of which could tolerate substitutions and account for species or subtype characteristics. HRH2 had a similar pattern of substitutions differentiating them from the other subtypes, while HRH3 and HRH4 had similar substitutions that align with their closer relationship. It was significant that none of the substitutions identified affected residues critical for ligand binding or the activation of the micro switch. Along with it, amino acid signatures, evolutionary distance, and Shannon entropy together indicate distinct biochemical and evolutionary features of histamine receptor subtypes. Leucine was predominant in HRH1–HRH3, but high alanine in HRH3 and high serine in HRH4 indicated subtype-specific patterns. On the evolutionary tree, HRH3 appeared as the most divergent subtype, HRH1 and HRH2 were relatively close together, and HRH4 was in the intermediate group. In the entropy maps, HRH1 showed the maximum variability and lesser variability from HRH2 to HRH4, suggesting enhanced evolutionary constraints. These studies together indicate a basic GPCR structure along with divergent evolution of subtypes for functional specialization.

Poly-string analysis indicated distinct subtype and species-specific repetition characteristics. HRH1 had a low number of repeats, with a conserved ‘WWW’ motif and some human-specific elements. HRH2 had even fewer repeats, in accordance with its extremely conserved sequence [42]. HRH3 had the highest degree of variation, with a species-specific glutamine heptamer and runs of alanine, methionine, and serine, in agreement with its specific compositional characteristics. HRH4 had only short and less frequent poly-strings, such as ‘SSS’ repeats, with a conserved ‘PP’ motif. Taken together, these indicate higher plasticity in HRH3 than HRH2 and HRH4. Intrinsically disordered analysis revealed characteristic subtype flexibility patterns. HRH1 displayed a higher degree of disordered and highly flexible residues, which reflect a more dynamic structure. HRH3 and HRH4 display moderate levels of disorder, whereas HRH2 displays the lowest levels, which reflect a generally rigid structure. These observations collectively imply that HRH1 uses its dynamic structure, involving flexible loops, to perform its function, whereas HRH2 retains a rigid structure that maintains its conserved role as a histamine receptor [60, 61]. The physicochemical Distance-based Phylogeny has revealed a subtype-specific grouping among Histamine Receptors. The most conserved group was formed by HRH3 and HRH4, while HRH1 demonstrated a moderate lineage-dependent dispersal. The subtype that was most internal but also most distant to other subtypes was HRH2, because it has a peculiar functional and physicochemical signature. The observed pattern of grouping is dominated by a subtype conservation and a clear subtype separation [62].

In summary, this extensive comparative analysis clearly illustrates that the mammalian histamine receptor family maintains a strongly conserved core structure of the GPCR family but follows distinct trajectories of subtype-specific evolution and diversification. The action of strong purifying pressures maintains the species-specific cores containing critical residues for ligand binding, activation, and G-protein coupling, whereas variations are selectively introduced into flexible and non-critical parts of the proteins to accomplish the fitness-related adaptations of the different subtypes by evolving a functional speciality or diversity.

## Author contributions statement

PG, SD, and AG initiated the research problem. SSH and DN developed the quantitative formulation. All authors contributed to executing the results and analyses. The primary draft was jointly written by all authors, who also reviewed and refined the manuscript to reach the final version.

